# TRF1 poly(ADP-ribosyl)ation by PARP1 allows proper telomere replication through helicase recruitment in non-ALT cells

**DOI:** 10.1101/2021.10.27.466083

**Authors:** C. Maresca, A. Dello Stritto, C. D’Angelo, E. Petti, E. Vertecchi, L. Pompili, F. Berardinelli, A. Sgura, A. Antoccia, G. Graziani, A. Biroccio, E. Salvati

**Affiliations:** Oncogenomics and Epigenetics Unit, IRCCS Regina Elena National Cancer Institute, Rome, Italy; Department of Biology and Biotechnology “Charles Darwin”, Sapienza University of Rome, Italy; Institute of Molecular Biology and Pathology, National Research Council, Rome, Italy; Department of Biology, Roma Tre University of Rome, Italy; Department of Systems Medicine, University of Rome Tor Vergata, Rome, Italy

## Abstract

Telomeres are nucleoprotein structures at eukaryotic chromosome termini. Their stability is preserved by a six-protein complex named shelterin. Among these, TRF1 binds telomere duplex and assists DNA replication with mechanisms only partly clarified. Poly (ADP-ribose) polymerase 1 (PARP1) is a chromatin associated enzyme which adds poly (ADP-ribose) polymers (PARs) to acceptor proteins by covalent hetero-modification. Here we found that TRF1 is covalently PARylated by PARP1 during DNA synthesis. PARP1 downregulation perturbs bromodeoxyuridine incorporation at telomeres in S-phase, triggering replication-dependent DNA damage and telomere fragility. PARylated TRF1 recruits WRN and BLM helicases in S-phase in a PARP1-dependent manner, probably through non-covalent PAR binding to solve secondary structures during telomere replication. ALT telomeres are less affected by PARP1 downregulation and are less sensitive to PARP inhibitors. This work unveils an unprecedented role for PARP1 as a “surveillant” of telomere replication, in absence of exogenous DNA insults, which orchestrates protein dynamics at proceeding replication fork.

## Introduction

Telomeres are nucleoprotein structures at eukaryotic chromosomes termini deputed to DNA end protection. They are non-genic regions consisting of specie-specific GC rich repeats bound by a six-members specialized complex called shelterin, which regulates telomere length homeostasis and prevents undesired recombination by repressing different pathways of DNA damage response[1][2]. Telomere duplication initiates from a single origin of replication, located at sub-telomeres, moving unidirectionally towards chromosome end. To this end, proceeding replication forks must cope with the compaction of telomeric heterochromatin and the presence of secondary structures (t-loops and G-quadruplex). Thus, telomere replication requires the action of several enzymes that are enriched at telomeric loci (helicases, topoisomerases, exonucleases, and ligases). Among these, the Telomere Repeat Binding Factor 1 and 2 (TRF1-2) were shown to facilitate the recruitment of RecQ helicases at telomeres [3][4]. TRF1 and TRF2 are members of the shelterin complex that directly bind to telomeric duplex, as homodimers, in a sequence-specific manner. Moreover, they interact and recruit other shelterins and chromatin remodeling enzymes to assist DNA replication and repair [2]. TRF1 loss has been shown to slow down replication fork progression at telomeres, consequently causing telomere fragility [5][6]. This effect is partially explained by the fact that TRF1 recruits the Bloom (BLM) RecQ helicase to replicating chromatin assisting DNA replication [4]. TRF2 has been shown to have a crucial role in pericentromeric chromatin replication, where it binds to SatIII satellite repeats and recruit[EP1]s topoisomerase I action [7].

PARP1 is the most abundant protein at chromatin after histones. It is responsible for the addition of poly(ADP-ribose) polymers (PAR) on proteins in response to DNA damage, but, as confirmed by several studies, poly(ADP-ribosyl)ation (PARylation) is also involved in various cellular pathways including transcription and chromatin organization. The immediate and robust PAR synthesis produced locally at damaged sites modifies protein-protein and protein-DNA interactions and serves as a molecular scaffold for the subsequent recruitment of chromatin modulators and DNA repair proteins [8]. PARP1 is in fact necessary to activate different DNA repair pathways and its inhibition induces synthetic lethality in the presence of functional defects of master regulator of DNA repair (i.e., BRCA2) [9]. At telomeres, PARP1 is implicated in DNA damage repair through activation of the alternative Non Homologous End Joining (alt-NHEJ) and Homologous Recombination (HR) [10]. Moreover, PARP1 interacts with and covalently modifies TRF2 [11]. Telomere specific PARPs (Tankyrase 1 and 2) are known to modify TRF1 and regulate telomere elongation and sister chromatids separation during mitosis. PARP1 is also enriched at telomeric chromatin during G-quadruplex stabilization, to resolve replication-dependent damage [12][13]. Here we investigate the constitutive role of PARP1 in difficult-to-replicate heterochromatin such as telomeric chromatin, in absence of DNA damage induction, unveiling a new role of this enzyme as a key modulator of protein dynamics at replicating telomeres.

## Results

### PARP1 and TRF1 interact in S-phase in a DNA independent manner

Telomeres require the shelterin protein TRF1 for the replication fork progression. In mice, TRF1 recruits the BLM helicase to assist DNA replication probably by removing secondary structures [4] in order to allow the replisome to passage. PARP1 cooperates with all five RecQ helicases to preserve genome integrity in replication stress conditions. [20] This led us to investigate if PARP1 and TRF1 could interact, and if this interaction could be implicated in DNA replication. To this aim, HeLa cells were synchronized at the G1-S boundary by double thymidine blockade. Cell cycle synchronization during progression into S and G2-M phases was measured by flow cytometry in the total cell population (Figure 1 A) and in BrdU pulsed cells to distinguish between early and mid/late S-phases of cell cycle, and the TRF1-PARP1 interaction was quantitatively assayed by co-immunoprecipitation at different time points after release (Figure 1 B). PARP1 was found immunoprecipitated by TRF1 and the affinity between the two proteins was found increased from the early S (time 0) to the mid-late S (2 hrs post release), while in G2-M (4 hrs post release) it returned to basal level (Figure 1 B). To ascertain whether PARP1 binding to TRF1 was dependent on the presence of DNA, co-immunoprecipitation experiments were performed in absence or in presence of ethidium bromide (EtBr, Figure 1 B). TRF1/PARP1 binding was strongly increased by EtBr addition, showing that this interaction did not require DNA; instead, TRF1/PARP1 interaction was increased in presence of EtBr, this could suggest that PARP1 had higher affinity for DNA-free TRF1, which abundance could be increased in presence of EtBr. To visualize a direct interaction between PARP1 and TRF1 *in-situ* in intact cells, PLA was performed (controls and experimental set up are shown in Figure S1), which revealed co-localization between proteins less than 40 nm far from each other, a distance at which two proteins are supposed to directly interact. PLA spots were detected in the nuclei of HeLa and analyzed by deconvolution microscopy (Figure 1 C). Signal quantification showed an increase in late S-phase cells, confirming the maximum of interaction during DNA replication (Figure 1 D).

**Figure 1:**
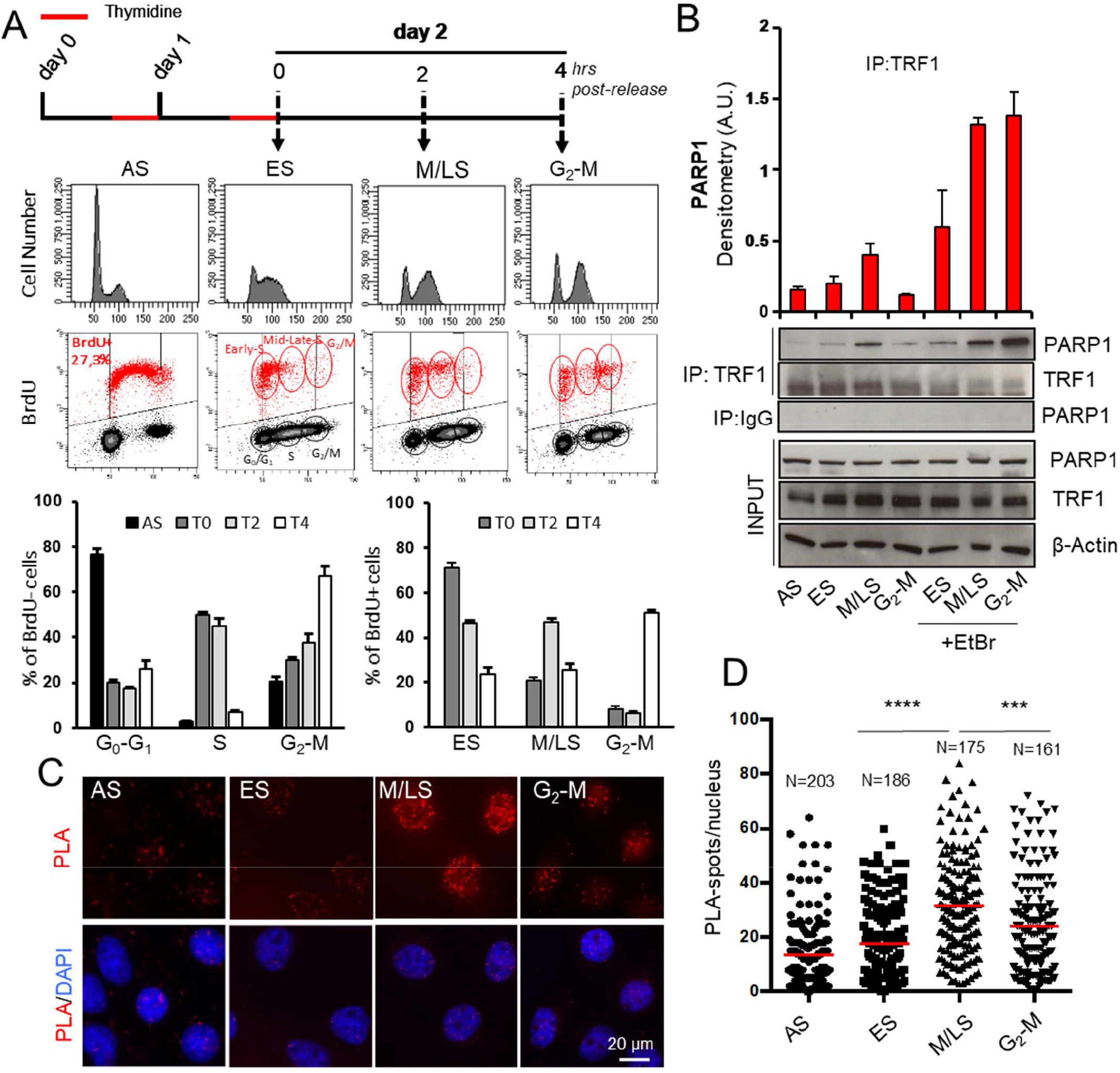
TRF1 and PARP1 interact during S-phase. HeLa cells were synchronized in the early S phase by double thymidine block and then released and collected at the indicated time points. A BrdU pulse was administered 15 minutes before the second thymidine block. Collected samples underwent cytofluorimetric analysis of the cell cycle phase distribution (A) or immunoprecipitation with an anti-TRF1 specific antibody or rabbit IgG as negative control (B) and decoration with the indicated antibodies (b-actin was used as loading control). Western blot signals were quantified by densitometry and reported in histograms after normalization on anti-PARP1 signals in the IgG immunoprecipitated samples, and anti-PARP1, TRF1 and b -actin signals in the input (B). One representative of three independent experiments is shown, bars are SD. C: HeLa cells synchronized as described were fixed in formaldehyde and processed for PLA with specific anti-TRF1 and PARP1 antibodies. Signals were acquired by Leica Deconvolution fluorescence microscope at 63X magnification (in C representative images are shown). The number of signals/nucleus was scored and reported in graph (D). For each column Mean and numerosity (N) are indicated, two pulled independent experiments were plotted, P value was determined by unpaired two tailed t-student test, *** P ≤ 0.001, **** P ≤ 0.0001

### TRF1 is covalently PARylated by PARP1 *in-vitro*

PARP1 synthetizes linear and branched PARs from NAD+ monomers, covalently linked to specific aminoacidic residues of PARP1 itself (homo-modification) or of specific target proteins (hetero-modification). To ascertain if TRF1 was directly modified by PARP1 enzyme, an *in-vitro* hetero-modification assay was performed in which recombinant TRF1 isoforms were added to PARP1 enzyme in presence of NAD+. The protein mixture was resolved onto PAGE and PARs covalently bound to PARP1 and TRF1 were detected by western blot analysis with an anti-PAR specific antibody (Figure 2 A). Full-length recombinant TRF1 was PARylated by PARP1 as shown by the appearance of a smear at a lower molecular weight in samples in which TRF1 was added (overlapping with the anti-TRF1 detected band shown in the right panel), compared to PARP1 signal alone. PARylation was further increased by cleaved DNA which stimulates PARP1 catalytic activity. TRF1 PARylation was also assessed by incorporation of biotinylated NAD+ in the Poly ADP-ribose polymers, in the heteromodification assay, after detection with HRP conjugated anti-Streptavidin. As shown in Figure 2B, the NAD+ incorporation is detected both at >100 KD (PARP1) and at 63KD (TRF1) when TRF1 is present, after biotin-NAD+ addition. This result, besides showing that TRF1 is a PARP1 substrate for covalent PARylation, further confirms an unprecedented direct interaction between the two proteins. In a non-covalent PARylation assay, recombinant TRF1 was immobilized onto a nitrocellulose membrane together with the H1 histone (a known PARP1 substrate of both covalent and non-covalent PARylation) and incubated with in vitro synthesized PARs, followed by anti-PAR detection. The dot-blot in Figure 2 C revealed that TRF1 is not a substrate for PARP1 non-covalent modification.

**Figure 2:**
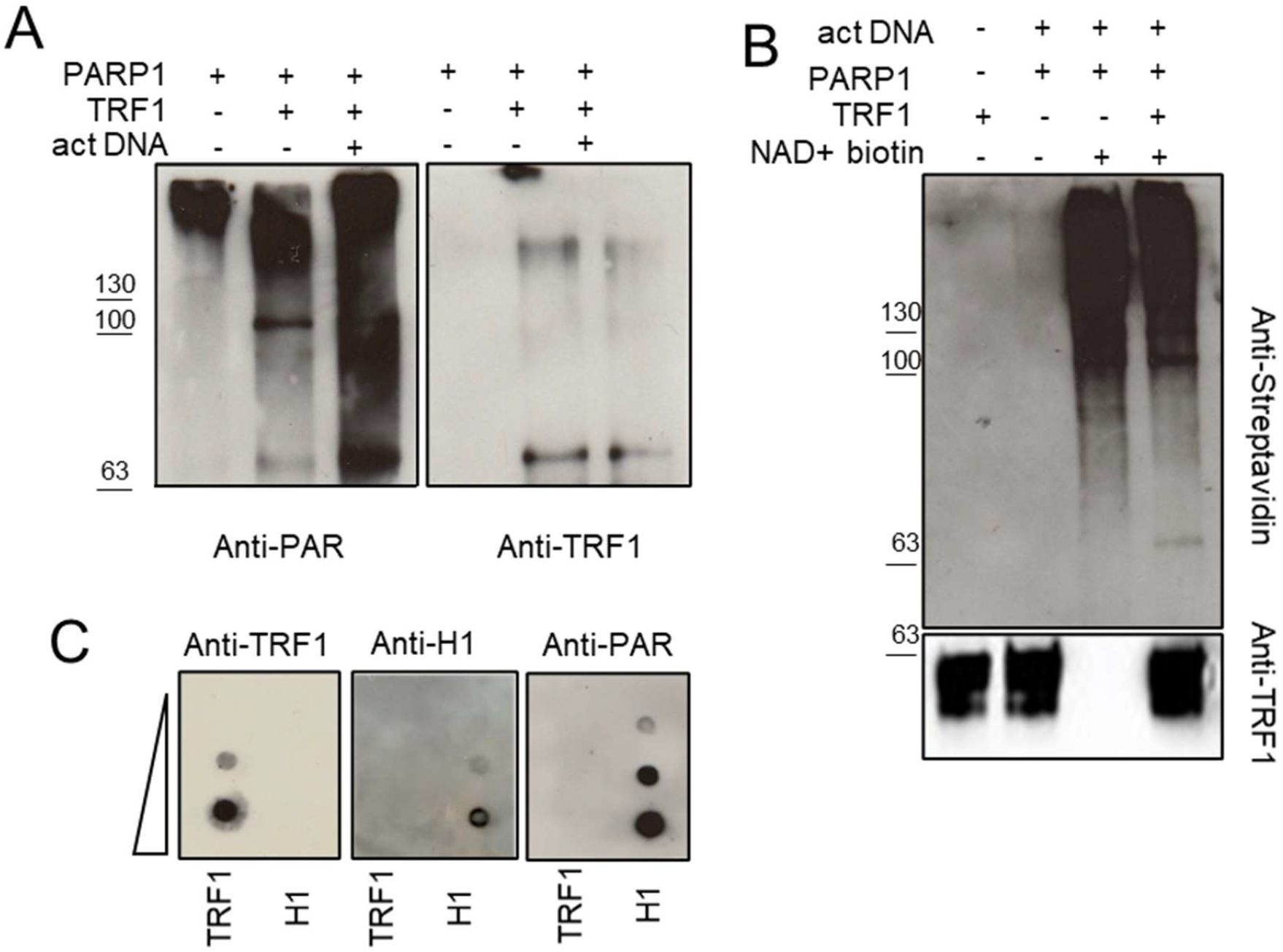
TRF1 is covalently PARylated by PARP1 *in-vitro*. **A**: high activity purified PARP1 enzyme was incubated with unlabeled NAD+ (**A** and **C**) or biotin-labelled NAD+ (**B**) in the PARylation reaction buffer in absence or presence of recombinant His-tag full length TRF1, with or without activating DNA (**A**). Protein mixtures were resolved on SDS-PAGE and incubated with an anti-PAR antibody (**A**) or HRP-Streptavidin (**B)** to detect PARylated proteins and anti-His or anti-TRF1 antibodies to detect TRF1 isoforms where present. Signals were revealed by chemiluminescence. **C**: Noncovalent PARylation assay: increasing quantities of recombinant full length TRF1 or H1 histone were spotted on nitrocellulose by dot blot, incubated with previously synthetized and purified PARs, and then decorated with anti-PAR antibody (to detect bound polymers), anti-TRF1 and anti-H1 antibodies and revealed by chemiluminescence.

TRF1 PARylation was finally detected in vivo in HeLa cells transfected with siTRF1 or control scrambled sequence (siSCR) and synchronized during progression through S-G_2_M phases of cell cycle. Samples collected at different time points underwent anti-PAR immunoprecipitation and detection with anti-TRF1 antibody. The anti-TRF1 blot in IP:PAR samples in Figure 3A clearly showed a band which increased during the late S-phase, following the same trend of TRF1/PARP1 affinity (in Figure 1), that was missing in siTRF1 interference (checked in the input samples) and IgG immunoprecipitated samples. PAR immunoprecipitation efficiency was controlled by incubating the entire gels of input and IPed samples with the anti-PAR antibody that showed an enrichment of PARylated proteins especially at low molecular weight (this is expected since histones are heavily PARylated). Interestingly, the immunoprecipitation with TRF1 and detection with anti-PAR in synchronized Hela cells, detected a band of the same molecular weight of TRF1 with a similar trend of accumulation through S-phase progression that was dependent on the presence of PARP1 protein (Figure 3B and C). The presence of a Tankyrase1 PARs acceptor site in the acidic domain of TRF1 was already shown [21]. However, in heteromodification assay, the delta acidic mutant of TRF1 was PARylated with the same extent of the full-length protein, demonstrating that PARP1 heteromodification engages other domains. As a control, PARylation of both full-length and delta acidic TRF1 variants was inhibited by the PARP1 inhibitor olaparib (Supplemental figure 2). Of note, as shown in Figure 3D and E, Tankyrase1 affinity for TRF1 during cell cycle, had an inverse trend with respect to PARP1 binding, showing a decrease during S and G2-M phase progression. This indicates a chronological and physical separation between TRF1/Tankyrase1 and PARP1/TRF1 complexes formation suggesting a functional difference between Tankyrase1- and PARP1-dependent TRF1 PARylation, according with the evidence that TRF1 is PARylated in-vivo in S-phase by PARP1.

**Figure 3:**
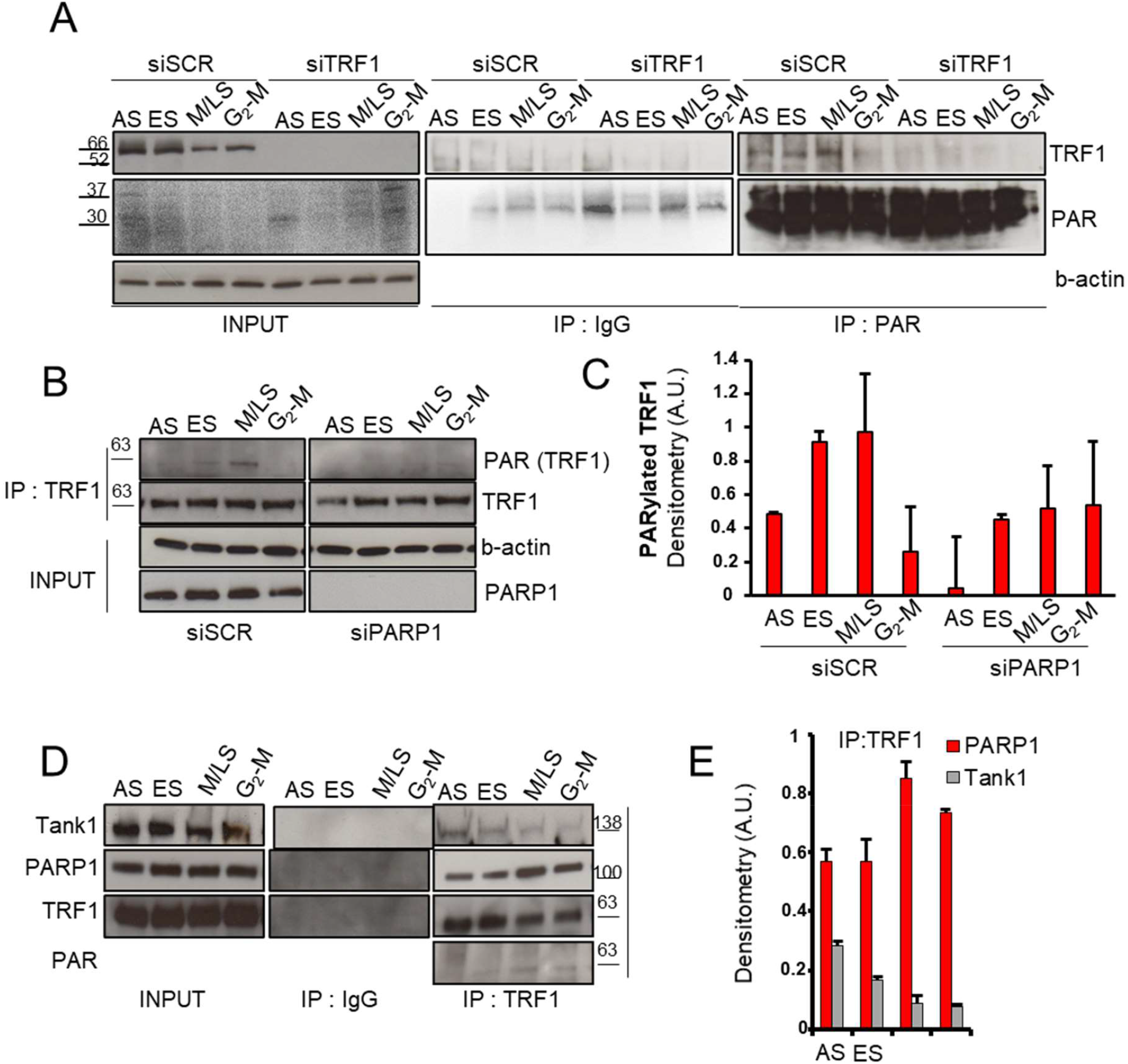
TRF1 is PARylated *in-vivo* in S-phase in a PARP1 dependent manner. HeLa cells were transfected where indicated, then synchronized, collected and immunoprecipitated as above described with the indicated antibodies or IgG as control. Input and immunoprecipitated samples were processed for Western blot analysis of the indicated antigens. **C**: quantification of **B, E**: quantification of **D**. One representative of three independent experiments is shown, bars are SD.

### TRF1 PARylation impacts on telomeric DNA replication

PARylation is known to alter the chemical environment of target proteins modifying their capacity to interact with other proteins and/or nucleic acids. It has been shown that TRF1 has a peculiar dynamic at telomeres during replication, detaching from chromatin during the replication fork passage[22]. Since PARP1 interacts and PARylates covalently TRF1 during S-phase, we wanted to ascertain if this interplay had a role in protein dynamics at replicating telomeres. To this aim, HeLa cells were interfered for PARP1 or a scrambled sequence and synchronized in the early S by double thymidine blockade. Then, 1 h before sample collection, cells were exposed to BrdU incorporation as indicated in Figure 4A. Samples were collected at different time points and splitted for different analysis. They were subjected to cell cycle distribution analysis (Figure 4 A), ChIP against TRF1 or BromoIP assays to analyze TRF1 association to telomeric chromatin and replication fork passage respectively (Figure 4 B). As shown in Figure 4 B, in control samples, TRF1 association to telomeric chromatin was reduced in the early S and in the G_2_-M phases, as expected, compared to non-synchronized cells. At the same time points, a peak of BrdU incorporation was observed, coherent with the model that TRF1 detaches from chromatin during fork passage, and with the fact that telomeres are replicated in two different times of cell cycle [22]. Interestingly, PARP1 interference delayed both the TRF1 dissociation and the BrdU incorporation (Figure 4 B). Since RNAi strategy could result in a hypomorphic phenotype, the same results were confirmed by using the PARP1 pharmacological inhibitors olaparib (Figure S3 A-C). At a dose unable to trigger DNA damage response activation (Figure S3 D), Olaparib treatment confirmed the lack of TRF1 dissociation and BrdU incorporation in the early S phase observed upon PARP1 interference. Since PARP1 depletion or inhibition seemed to impair TRF1 dissociation from chromatin, we deeper investigated the interplay between TRF1/PARP1 and telomeric duplex DNA. In the Electro Mobility Shift Assay shown in Figure 4 D, unmodified TRF1 efficiently bound ^32^P-labelled telomeric duplex DNA, but the binding was massively decreased by the previous heteromodification of TRF1 by PARP1 enzyme. As a control, PARP1 alone did not affect DNA migration. Although FACS analysis in Figure 4 A failed to reveal differences in cell cycle distribution between control and PARP1 interfered population, a more accurate analysis of S-phase length by the BrdU pulse experiment in Figure 4 C clearly shows that PARP1 interfered cells incorporated less BrdU and had a delayed S-phase exit. This is coherent with a localized impairment of DNA synthesis able to slow down S-phase exit but without effect on the whole population cell cycle distribution.

**Figure 4:**
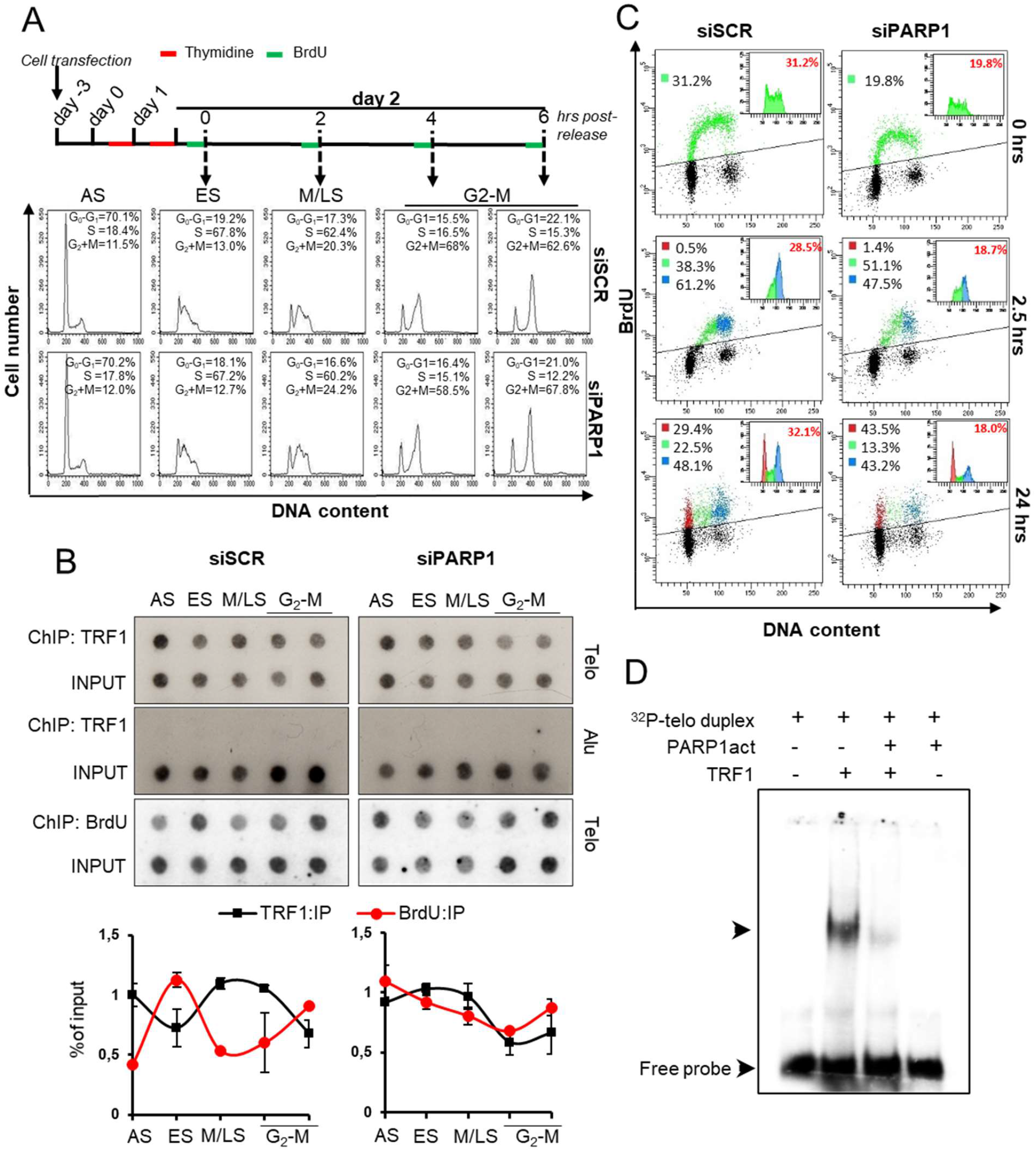
PARP1 inhibition perturbs DNA synthesis and TRF1 dynamics at telomeres in S-phase. HeLa cells were transfected with siSCR and siPARP1 RNAs, synchronized by double thymidine block as above described and one hour before collection were exposed to BrdU incorporation (**A**). Samples were collected, splitted and processed for FACS analysis (**A**), one representative of three independent experiments is shown, and ChIP against TRF1 or BrdU IP (**B**). Immunoprecipitated chromatin was dot blotted and hybridized with a radiolabeled probe against telomere repeats or Alu repeats (**B**) One representative of three independent experiments with similar results is shown. Immunoprecipitated samples signals were quantified by densitometry, normalized on each relative input (1:100) and Alu signal (where present) and then reported as the percentage of immunoprecipitated chromatin (**B**). Curves report the mean of three independent experiments, bars are SD. **C**: Bivariate distributions (dot plot) of BrdU (Alexa Fluor 488) content versus DNA (PI) content were analyzed. HeLa cells interfered as above described were pulsed with BrdU for 15 min, and after the indicated intervals in BrdU-free medium the DNA was denatured, incubated with anti-BrdU antibody and staining DNA. BrdU− (black area) and BrdU+ (multicolor area) populations were separated by analytical sorter in bi-parametric distribution and graph insert on top-right show DNA content of BrdU-positive cells, the percentage of positivity at each time point is reported in red inside the box. Flow cytometry data analysis is built upon the principle of gating and the percentages of G0-G1 (red) S (green) and G2+M (blue) was reported inside the dot plot. One representative of three independent experiments with similar results is shown. **D**: EMSA assay, radiolabelled DNA was incubated with unmodified or PARP1 PARylated TRF1 and run on nondenaturing polyacrylamide gel. Signals were acquired at the Phosphoimager.

### PARP1 inhibition induces transient DNA damage in telomerase positive cells but not in ALT cells

The impairment of replication fork progression at telomeres is expected to give rise to a transient activation of DNA damage response (DDR), revealed by the activation of γH2AX foci, due to the presence of single stranded DNA lesions in proximity to paused or stalled replication forks. Therefore, the expression of the above marker was analyzed at different time points after PARP1 down-regulation (via RNAi) in comparison with TRF1 down-regulation, as a control of telomere replication perturbation, in both HeLa and U2OS cell lines, the first with telomerase activity and the last adopting alternative telomere elongation mechanisms (ALT) involving the break-induced DNA synthesis. The effect of the double interferences was also analyzed to ascertain whether PARP1 and TRF1 were acting in the same pathway (Figure 5 A-C). The single cell analysis by immunofluorescence-FISH co-staining of telomeric DNA and phosphorylated γH2AX, showed a transient increase of the percentage of γH2AX positive cells (Figure S4) and of TIFs (recognized as γH2AX/telomere colocalizations) positive nuclei in both TRF1 and PARP1 interfered samples in HeLa cells that were recovered at 72 h after interference. In addition, in double interfered samples, TIFs positive cell percentages were like the TRF1 single interfered samples, indicating that PARP1 and TRF1 acts epistatically (Figure 5 A and B and Figure S4 A and B). Interestingly, in U2OS cells, the PARP1 interference was almost ineffective both alone and in combination with siTRF1 (Figure S4 A and C and Figure 5 A and C). The extent of protein down-regulation achieved by interference is shown in Figure S4D.

**Figure 5:**
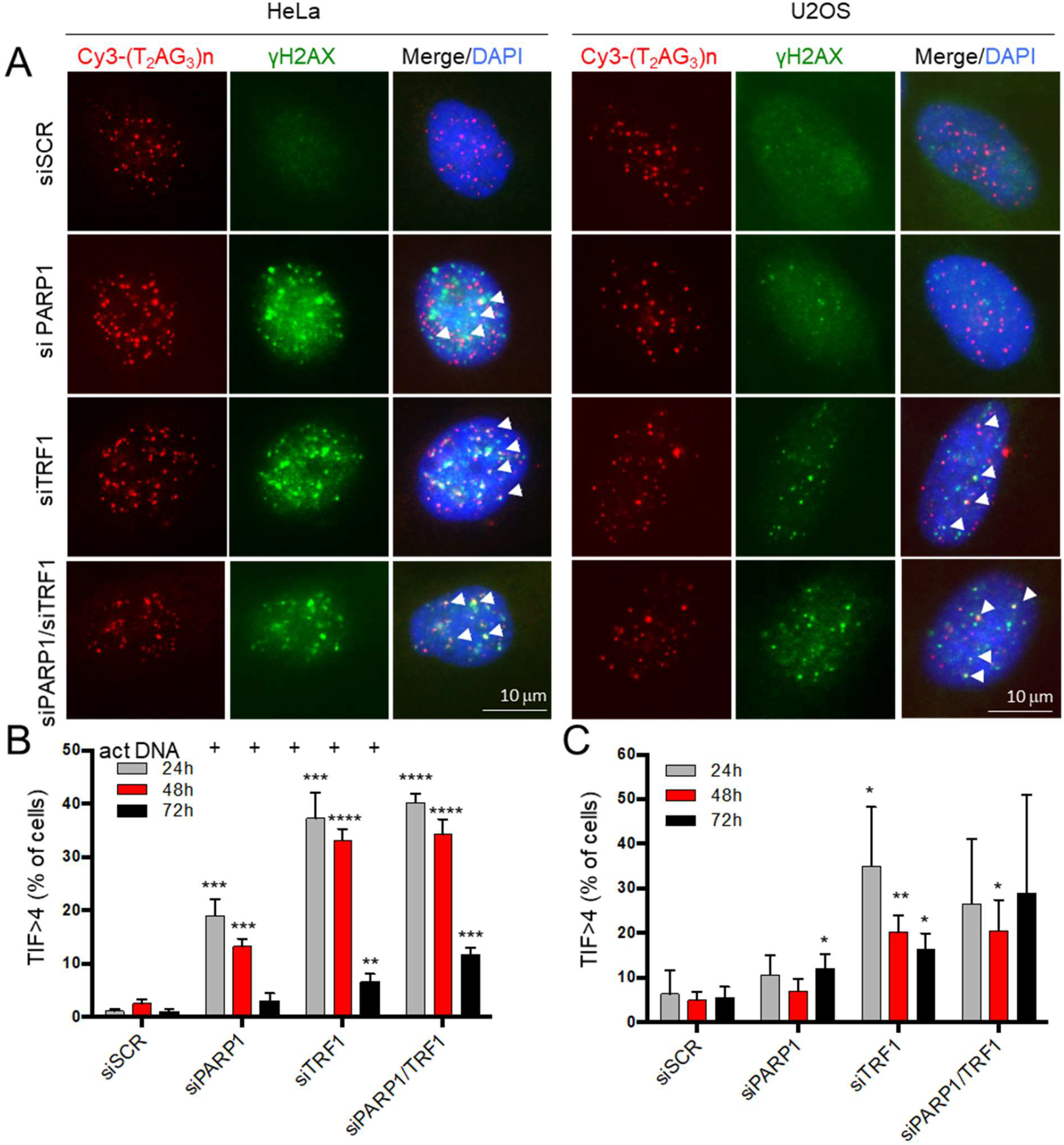
PARP1 inhibition causes transient DNA damage at telomeres in non-ALT cells. HeLa and U2OS cells were transfected with siRNAs against siPARP1 and siTRF1 alone and in combination and against a scrambled sequence. Then samples were fixed at the indicated endpoints after transfection and processed for IF-FISH against gH2AX and telomere repeats with a Cy3-Telo PNA probe and counterstained with DAPI. Signals were acquired by Leica Deconvolution fluorescence microscope at 63X magnification, representative images are shown in panel A. The percentage of TIFs positive cells (displaying >4 g H2AX/telomere co-localizations) in HeLa and U2OS cells was scored and reported in histograms in B and C, respectively. The average of three independent experiments is shown, bars are SD. Bars are SD. *P≤0.05, ** P ≤ 0.01, *** P ≤ 0.001, **** P ≤ 0.0001.

### PARP1 inhibition causes replication-dependent DNA damage and telomeric fragile sites in non-ALT cells by interfering with helicase recruitment

The transient nature of telomeric damage observed upon PARP1 inhibition/interference underlies the activation of DNA repair to resolve fork pausing/stalling. To ascertain if the DNA damage induction observed upon PARP1 interference was due to replication perturbation, we analyzed the activation of pRPA(S4/S8), which is considered as a marker of forks collapse, at telomeres. The percentage of pRPA/telomere colocalization positive cells increased in both TRF1 and PARP1 depleted cells, indicating that telomeric DNA damage was due to replication defects undergoing repair, and, of note, the double interference failed to show further increase (Figure 6 A and B). Replication stress at telomeres is known to generate a phenotype of telomere fragility, recognizable by the presence of double telomeric spot at a single chromatid in telo-FISH assay. Consistently with this finding, PARP1 interference (Figure 6 C and D), as well as pharmacological inhibition (Figure S3 E-F), were able to induce a significant increase of telomere fragile sites, comparable to the TRF1 interference. Also, in this case, the double TRF1/PARP1 knock-down displayed a similar result compared to the PARP1 knock-down. Neither TRF1, nor PARP1 or their combination affected telomere length (Figure 6E and S5 A). Other kind of telomere aberrations, not related to telomere replication (i.e. telomere fusions and dicentric chromosomes) were scored without finding any significant increase with respect to control sample (Figure S5 B).

**Figure 6:**
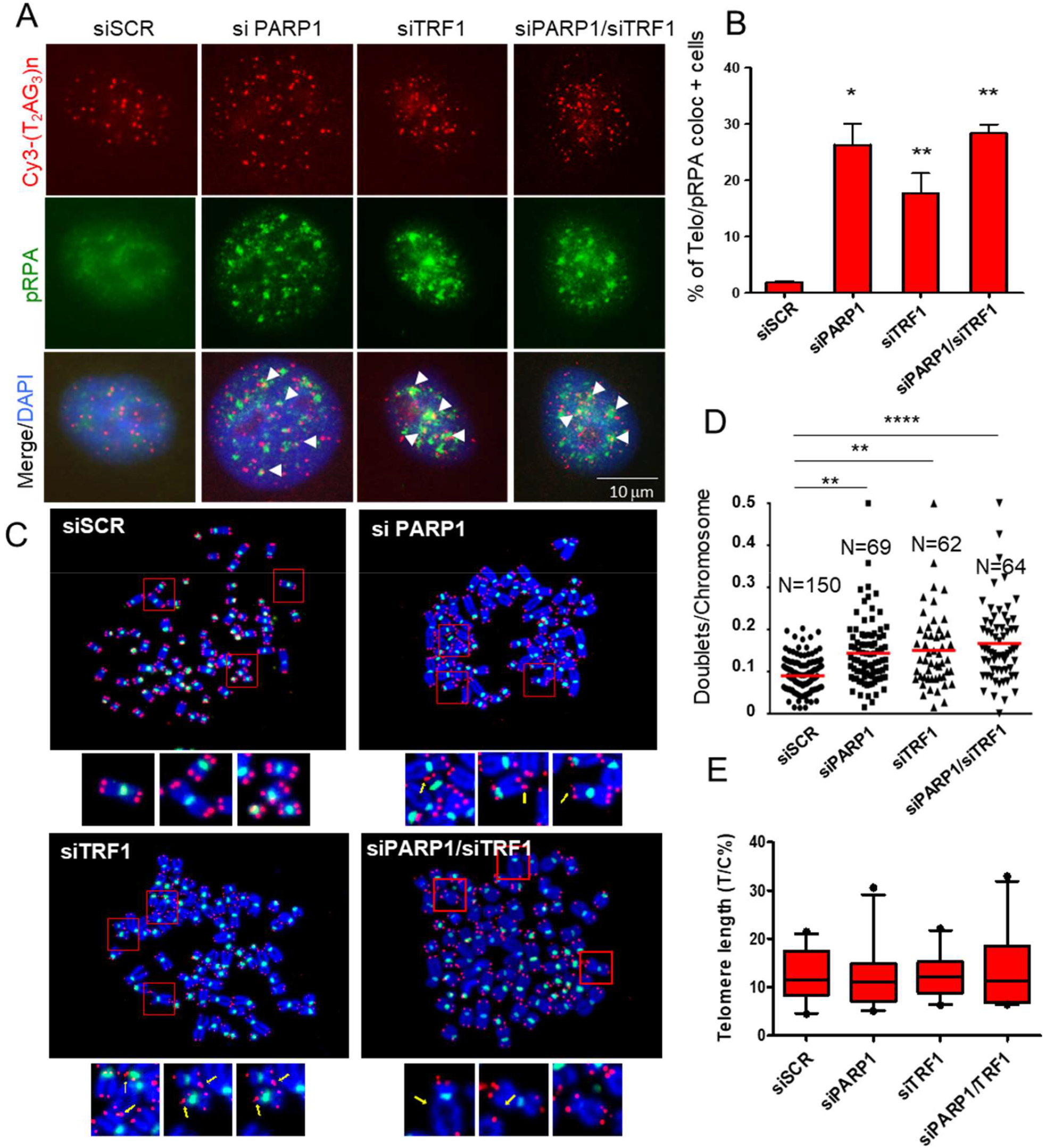
PARP1 inhibition induces telomere replication defects. HeLa cells were interfered and fixed as above indicated, and then processed for IF-FISH against pRPA and telomere repeats with a Cy3-Telo PNA probe and counterstained with DAPI. Signals were acquired by Leica Deconvolution fluorescence microscope at 63X magnification (representative images are shown in panel **A**). Telomere/pRPA colocalizations were scored and the percentage of positive cells (displaying >4 colocalizations) was reported in histograms (**B**). HeLa were transfected as above and after 72 hours metaphases were collected and processed for fish for pantelomeric/pancentromeric staining and counterstained with DAPI. Representative images at 100X magnification are shown in **C. D**: Telomere doublets were scored and reported in graphs as the percentage of doublets/chromosomes Two pulled independent experiments were plotted; **E**: Q-FISH analysis of telomere length on the same metaphases. P value was determined by unpaired two tailed t-student test. ** P ≤ 0.01, **** P ≤ 0.0001.

In agreement with the lack of TIFs induction upon siTRF1 and siPARP1 interference, U2OS cells were also resistant to the induction of telomere fragility (Figure 7A and B). Interestingly, pharmacological PARP1 inhibition by olaparib had different effects on HeLa and U2OS cells which displayed a significantly different IC50 to the PARPi (Figure S9). By extending the analysis to other cell lines of different histological origin, previously characterized for the presence of ALT mechanisms, we discovered that ALT cells displayed lower sensitivity to olaparib cytotoxic effects (Figure 7 C and Figure S9).

**Figure 7:**
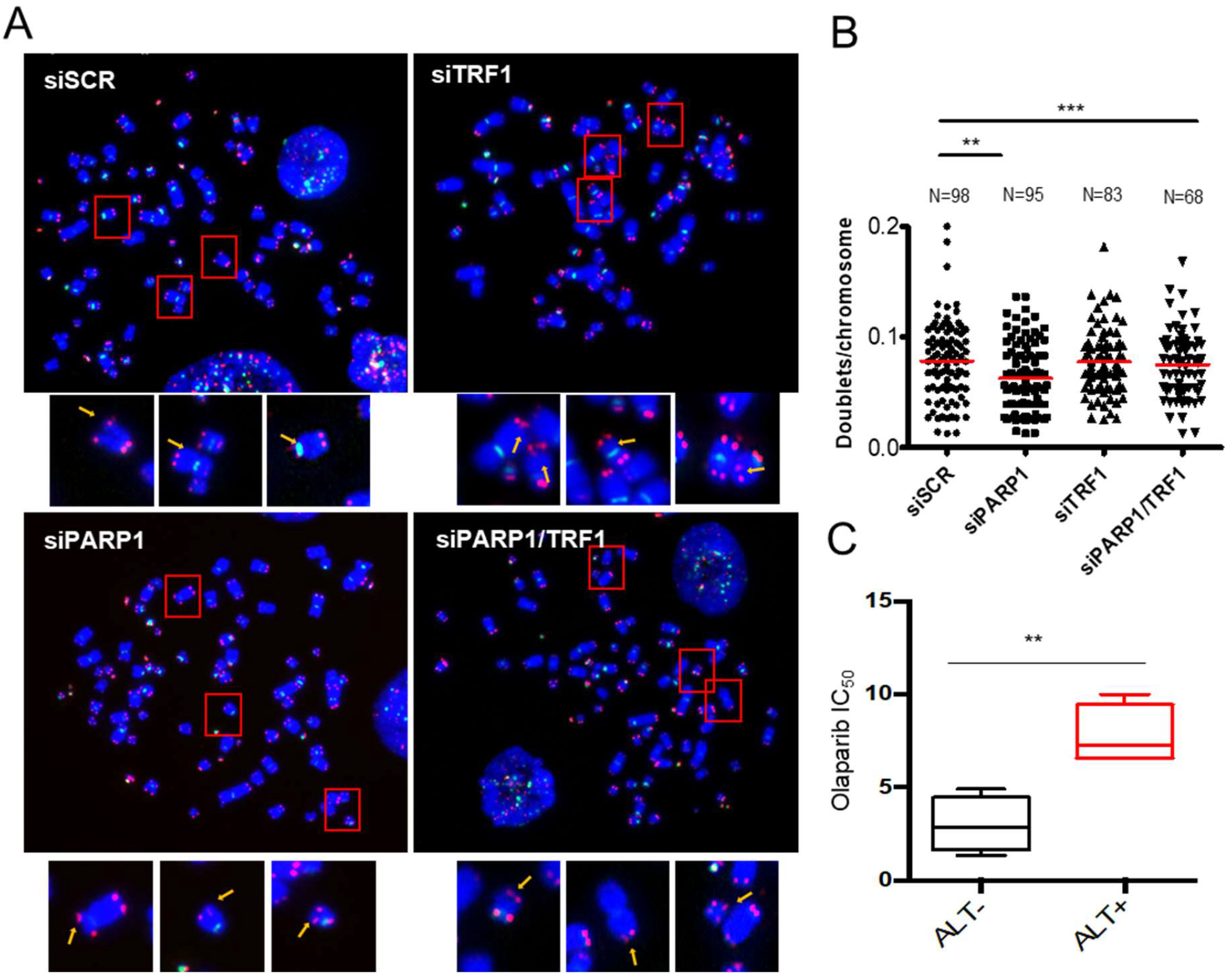
ALT cells are resistant to fragility induced by siTRF1 and siPARP1 and are less sensitive to Olaparib treatment. U2OS cells were transfected with siRNAs against siPARP1 and siTRF1 alone and in combination and against a scrambled sequence and after 72 hours metaphases were collected and processed for FISH for pantelomeric/pancentromeric staining and counterstained with DAPI. Representative images at 100X magnification are shown in A. B: Telomere doublets were scored and reported in graphs as the percentage of doublets/chromosomes. Two pulled independent experiments were plotted, bars are means; C: ALT and non-ALT cell lines (4 cell lines for each group) were exposed to olaparib concentrations ranging from 0.5 to 10 mM for 7 days. Cell survival was determined by crystal violet and IC50 was calculated and reported in boxplots. Bars are SD. *P≤0.05, ** P ≤ 0.01, *** P ≤ 0.001, **** P ≤ 0.0001.

The mechanisms underlying telomere doublets formation is not completely clarified. It was already shown that TRF1 recruits the activity of BLM RecQ helicase to resolve topological stress at replicating telomeres and the lack of BLM recruitment by TRF1 is responsible for telomere fragility phenotype [4]. Here we show that TRF1 co-immunoprecipitated also with WRN, another RecQ helicase (Figure 8 A and B). More interestingly, both the RecQ helicases are recruited by TRF1 in a PARP1 dependent manner in S-phase (Figure 8 A and B). ChIP analysis of WRN association to telomeric chromatin during S-phase showed a peak in early S-phase, that was completely abrogated in PARP1 interfered cells (Figure 8 C and D). The lack of WRN recruitment explains the inability of cells with downregulated PARP1 to complete telomere replication and the formation of fragile sites.

**Figure 8:**
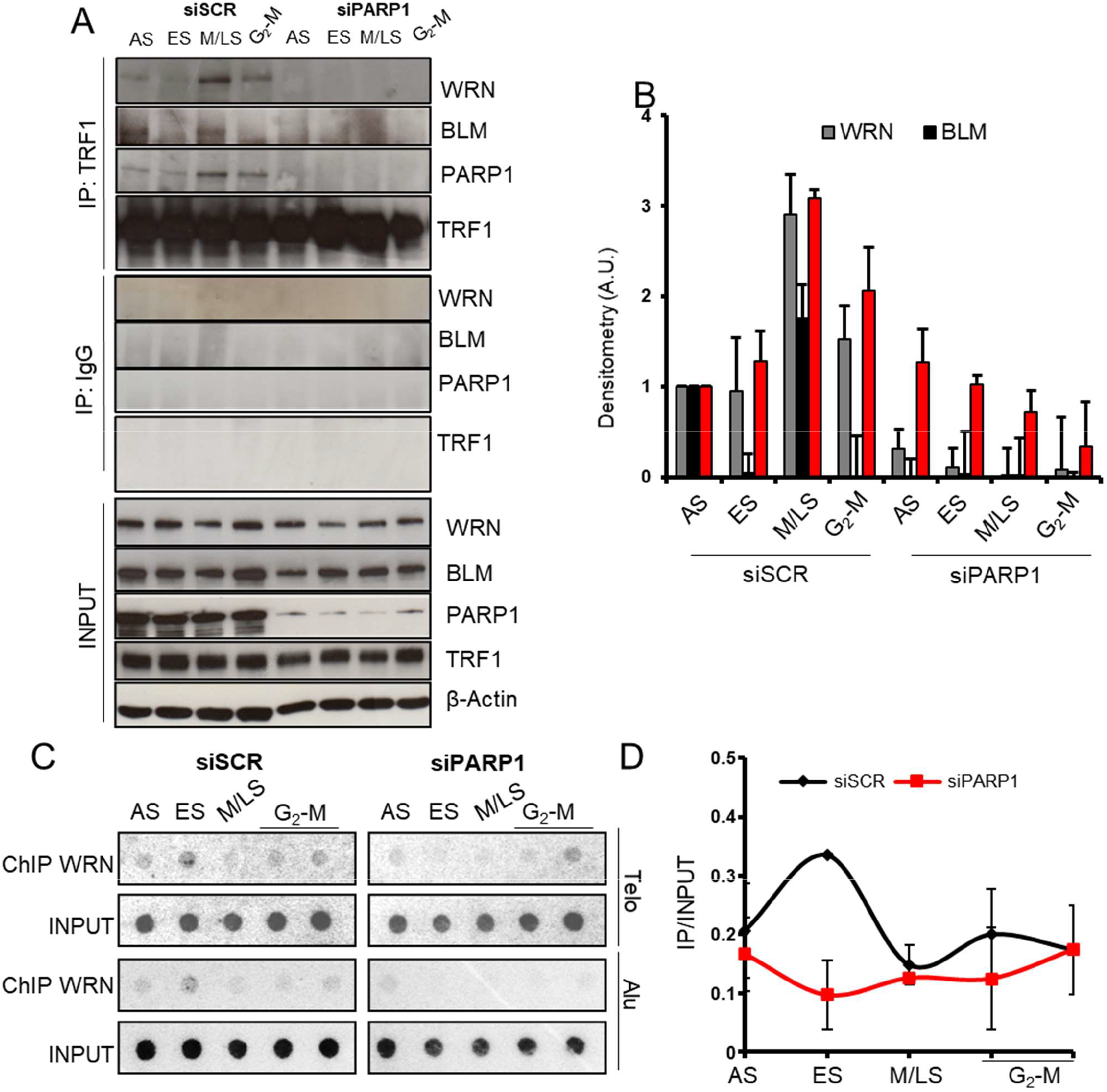
PARP1 inhibition impairs WRN and BLM recruitment by TRF1 during S-phase. **A**, HeLa cells interfered for PARP1 and synchronized as above reported were immunoprecipitated against TRF1 and decorated for the indicated antigens. Densitometry of immunoprecipitated proteins, normalized on each respective input is reported as histograms (**B**). Samples processed as in **A** underwent ChIP against WRN. Immunoprecipitated chromatin was dot blotted and hybridized with a radiolabeled probe against telomere repeats or Alu repeats (**C**) Immunoprecipitated samples signals were quantified by densitometry, normalized on each relative input, Alu signal was subtracted to each relative sample and then reported on graphs as IP/Input ratio (**D**). One representative of three independent experiments with similar results is shown. Graphs report the mean of three independent experiments, bars are SD.

## Discussion

Here we unveil an unprecedented interaction between the shelterin protein TRF1 and the PARP1 enzyme. PARP1 is canonically activated by DNA damage and can synthetize PAR chains on itself and on specific acceptor proteins by modifying the molecular environment around the DNA lesion facilitating repair actions. In this paper, we observed a specific function of PARP1 at telomeres during DNA synthesis that is functional to proper DNA replication. At first, we assessed a direct interaction between PARP1 and TRF1, which does not require the DNA presence and is S-phase dependent (Figure 1). In addition, we discovered that TRF1 is a substrate for PARP1-dependent covalent PARylation but is unable to bind PARs through non-covalent interaction (Figure 2). This observation allows to exclude that TRF1 could interact with auto-PARylated PARP1, sustaining the hypothesis that TRF1 is a specific PARP1 target. The functional role of PARP1 and PARylation during telomere replication was assessed by the Bromo-ChIP experiments in which it is clearly demonstrated that both the lack of the protein and the catalytic inhibition, achieved with two different PARP1 inhibitors, impairTRF1 dynamics and BrdU incorporation (Figure 3, Figure S4 and Figure S5). Since PARylation is known to add negative charges to target proteins altering protein-DNA affinity, we hypothesized that TRF1 PARylation decreases TRF1 binding to DNA duplex facilitating the access of the replisome. In agreement with this finding, the EMSA assay confirmed that PARylation by PARP1 impairs TRF1 binding to telomeric duplex (Figure 4D). Moreover, since TRF1/PARP1 affinity was increased by DNA degradation (in presence of EtBr, Figure 1 B), we could infer that, during the fork passage, PARylated TRF1, displaced from DNA duplex, can form multiprotein complexes recruiting WRN and BLM helicases to unwind secondary structures such as G-quadruplex formed on the lagging strand of the proceeding fork. Both BLM and WRN are covalent and non-covalent PARs binders cooperating with PARP1 in maintaining DNA integrity [20]. This means that they can interact with PARP1 via protein-protein interaction, or with PARylated substrates via non-covalent interaction (here TRF1). This suggests a model in which PARP1, while binding and PARylating TRF1 to remove it from the telomeric duplex during fork passage, also allows recruitment of both BLM and WRN, by protein-protein interaction or by PARs-mediated interaction, to remove secondary structures forming on the G-rich lagging strand (Figure 9). In agreement with this, PARP1 interference completely abrogates the recruitment of WRN and BLM by TRF1 during S-phase (Figure 8 A and 8 B). The consequence of the lack of helicase recruitment generates a replication dependent DNA damage confirmed by the transient activation of γH2AX and RPA phosphorylation, which indicates the presence of a single stranded DNA lesion due to ongoing DNA repair. The cell cycle distribution is overall unaffected in PARP1 interfered cells, suggesting that PARP1 inhibition does not affect the whole DNA synthesis. However, the S-phase length analysis, performed by a BrdU pulse incorporation, evidenced a delay of S-phase that is coherent with a perturbation of DNA synthesis localized at telomeres. Of note, DDR activation is not visible in U2OS upon PARP1 interference. Surprisingly, while siTRF1 is still able to induce DDR in U2OS, both TRF1 and PARP1 depletion are not effective in inducing telomere fragility (Figure 6 and 7). U2OS are known to activate alternative mechanisms of telomere length maintenance involving break-induced replication (BIR). ALT cells are characterized by long and heterogeneous telomeres with different epigenetic structure. ALT cells display high replication stress at telomeres, which triggers frequent recombination and telomere elongation through different mechanisms [23]. It has been recently reported that ALT cells display telomere fragility at least in part caused by BIR events, in which BLM takes part, and that alt-Non Homologous End Joining suppresses BIR and telomere fragility in non-ALT cells [24]. Nevertheless, here we found that neither TRF1 nor PARP1 depletion led to the formation of fragile telomeres in ALT cells, unveiling new levels of complexity for these mechanisms.

**Figure 9:**
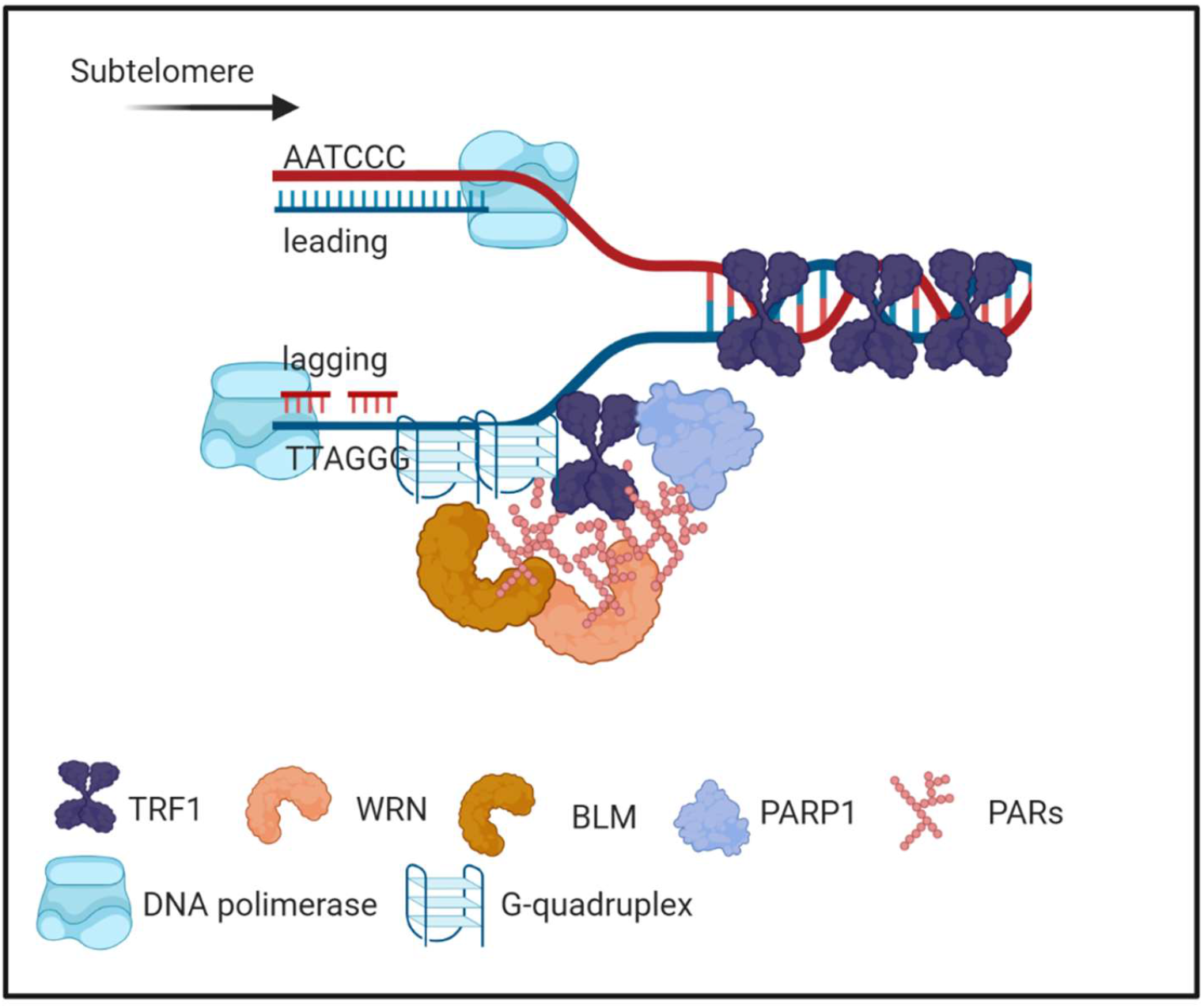
Model for TRF1/PARP1/RecQ helicases interplay during telomere replication. PARP1 PARylated TRF1 has lower affinity for DNA duplex and is recruited in a complex with BLM and WRN that resolve secondary structures forming at the G-rich lagging strand. Created in BioRender.com.

Other authors reported that a stable PARP1 knock-out triggered loss of telomere repeats and DDR induction in colon cancer cells [25]. Apart of telomere doublets, here we did not observe a reduction of telomere length or an increase of other telomeric defects (Figure 6 and S8), but this could be explained by the fact that we analyzed cells in the first round of duplication after a transient knock-down. In addition, the double TRF1/PARP1 interference had the same effect of the single ones indicating that TRF1 knock-down phenotype is recapitulated by PARP1 inhibition. This evidence strongly supports the idea that helicase recruitment is dependent on PARP1 and/or PARylation, at least in telomerase positive cells, and is at the basis of the telomere replication defects observed upon TRF1 knock-down. Terminal forks are supposed to frequently pause and stall, especially in actively replicating cells, as tumor cells are. We can reasonably suppose that a PARP1 “surveillance” activity is required during telomeric DNA synthesis.

In conclusion, this work provides mechanistic insights on how PARP1 orchestrates the molecular events occurring at replicating telomeres through covalent modification of TRF1 and recruitment of BLM and WRN helicases. The PARP1 “surveillance” seems to be specific for non-ALT cells, which display less replication stress and more stable genotypes suggesting that the PARP1 “surveillance” could be lost in cancer evolution along with the increase of genetic instability.

## Materials and methods

### Cell cultures, transfections, and treatments

Human cervical cancer cells (HeLa), BJ human fibroblasts and human osteosarcoma cells (U2OS) were purchased from ATCC repository. BJ EHLT were obtained by retroiviral transduction of BJ cells with hTERT (Addgene plasmid #1773) and Large T SV40 antigen (Addgene plasmid # 21826) as described. Cells were maintained in Dulbecco Modified Eagle Medium supplemented with 10% fetal calf serum, 2 mM L-glutamine, and antibiotics. In synchronization experiments, cells were seeded at 40% confluence and exposed to 2 mM thymidine (T-1895, SIGMA) for 16 h (I block), followed by 8 h release in fresh medium, and again exposed to 2 mM thymidine for additional 16 h (II block). Then cells were released in fresh medium and collected by trypsinization at different time points for further analysis. RNAi was performed by transfecting cells 2 days before synchronization at 20% confluence with 5 nM siRNA (scrambled sequence, two different sequences against PARP1 and TRF1: PARP1 siRNA Origene SR300098B/C, TRF1 siRNA Origene SR322000B/C, SCR siRNA Origene SR30004 and POLYPLUS INTERFERIN #409-10 as Transfection reagent).

### Flow cytometry

Cell cycle analysis was performed by flow cytometry (Becton-Dickinson) after cellular staining with propidium iodide (PI), as previously described [14][15]. After culturing and treatment, cells were harvested, washed with PBS twice, fixed in 70% ethanol at 4 °C overnight. Then, cells were washed with PBS twice, stained with PI at a final concentration of 50 μg/mL and RNase at a final concentration of 75 kU/mL, incubated for 30 min, then analyzed by FACSCalibur and FACSCelesta (BD Biosciences, San Jose, CA, USA). Progression of cells through the cell cycle phases was analyzed by simultaneous flow cytometric measurements of DNA and 5-bromo-2′-deoxyuridine (BrdU) contents of cells, as previously described (Biroccio et al. 2001). Briefly, cells were pulsed with BrdU (Sigma Aldrich) at a final concentration of 20 μM for 15 min, and after the appropriate intervals in BrdU-free medium (from 2.5 to 24 h) the DNA was denatured. Cells were then incubated with 20 μl of the mouse Mab-BrdU (347580 Pure BD) for 1 h at room temperature, and BrdU-labeled cells were detected using goat anti-Mouse Fab′2 Alexa Fluor 488 (Cell Signaling). The cells were counterstained with PI, acquired and analyzed with BD FACS Diva Software.

### Immunoprecipitation and western blot

Cells treated as above were collected and lysed in nuclei isolation buffer (10 mM Hepes pH 7.5, 10 mM KCl, 0.1 EDTA, 0.1 mM EGTA, 0.1 mM DTT, protease and phosphatase inhibitors). Nuclei were isolated by centrifugation and lysed in high salt RIPA buffer (50 mM Tris-HCl ph7.4, 330 mM NaCl, protease and phosphatase inhibitors). For immunporecipitation, 500 μg of proteins were incubated with 4 μg of goat IgG, anti-TRF2 (Mouse Mab Millipore 05521), anti-TRF1 antibody (Goat Pab sc-1977, SantaCruz), or anti-PAR (Mouse Mab 10H ALX-804220, Alexis) recovered with Protein-G dynabeads (Invitrogen), run on PAGE together with input sample (1:20 of amount of immunoprecipitated proteins) and blotted with anti-PARP1 (Mouse Mab 551025 BD Pharmigen), anti-TRF1 (Rabbit Pab sc-6165, Santa Cruz), anti-Tankyrase1 (Mouse Mab IMEGENEX IMG-146), or anti-PAR (mouse Mab 10H ALX-804220, Alexis); β-actin was used as a loading control (mouse Mab Sigma A2228). Five U/μg DNA of DNase (Roche) or ethidium bromide (1%) were added in the lysate before immunoprecipitation with anti-TRF1 antibody (goat Pab sc-1977, SantaCruz).

### Protein expression and purification

His-tagged human wild-type (wt) and delta-acidic TRF1 were expressed in Escherichia coli BL21(DE3) by using pTrc-HisB vectors (a kind gift of Prof. Eric Gilson, University of Nice-Sophia Antipolis, Nice). TRF1 mutants expression was induced at an OD 600 of 0.3–0.4 with 1 mM IPTG, followed by an incubation for 4 h at 37°C. After centrifugation, cells were resuspended in lysis buffer [50 mM sodium phosphate pH 7.2, 300 mM NaCl, 10 mM imidazole, 1 mg/ml lysozyme, PMSF]. Cells were sonicated and the insoluble fraction was removed by centrifugation at 15,000 g for 30 min. The soluble fraction was loaded on 1.5 ml of HisPur™ Ni-NTA Resin (Thermo Fisher scientific Inc.) and incubated 1 hour at 4°C on rotation. Elution was performed with 250 mM imidazole in a buffer consisting of 50 mM sodium phosphate pH 7.2, 300 mM NaCl. Elution fraction was run on PAGE and quantified by Coomassie staining.

### Heteromodification of HIS-hTRF-1 isoforms by PARP1

For the analysis of TRF1 PARylation by PARP-1 [16], beads were pelleted (160 ng of hTRF-1/sample) and re-suspended in activity buffer containing 5 units of hPARP-1 (High Specific Activity, Trevigen), 2.5 μg DNase I-activated calf thymus DNA, 200 mM NAD+, Tris-HCl pH 8, 10 mM MgCl_2_ and 2 mM dithiothreitol. In control samples hPARP-1 or hTRF-1 were omitted. Reaction specificity was evaluated by adding to the reaction mixture 1 mM of the PARP inhibitor olaparib (Selleckchem). As positive control of PAR covalently bound to an acceptor protein, 10 μg of histones (Merck Millipore) were added to the reaction mixture containing PARP-1. After 30 min of incubation at 25°C, the reaction was stopped by the addition of Laemmli sample buffer and samples were analyzed by gel electrophoresis on 8% SDS-PAGE and Western blot. PARylated PARP-1, TRF-1 or histones were detected using anti-PAR monoclonal antibody (Trevigen) and input TRF1 mutants were revealed with anti-His (anti 6-His Rabbit Pab Sigma Aldrich). For the detection of biotin-labelled PARylated proteins the same assay was conducted in presence of biotin-NAD+ (Sigma Aldrich) followed by SDS-PAGE and western blot detection with anti-streptavidin HRP antibody (Molecular Probes).

### Synthesis of PAR and non-covalent PAR binding

Synthesis of PAR was performed as previously reported [17]. Briefly, 50 units of purified human PARP-1 (High Specific Activity hPARP-1, Trevigen) were incubated in a mixture containing 100 mM Tris-HCl pH 8, 10 mM MgCl_2_, 2 mM dithiothreitol, 2.5 μg of DNase I-activated calf thymus DNA (Trevigen) and 200 mM NAD+ (Sigma-Aldrich) for 45 min at 30°C. The reaction was stopped by adding ice-cold trichloroacetic acid (TCA) to a final concentration of 20% (w/v). PARs were detached from proteins by incubation in 50 mM NaOH and 10 mM EDTA for 1 h at 60°C. After adjustment of pH to 7.5, PAR were purified by phenol/chloroform extraction as described [17]. For the study of non-covalent interaction of PAR with TRF-1, graded concentrations of purified His-hTRF1 protein were immobilized directly by slot-blotting on nitrocellulose membrane. Histone H1 (Millipore) was used as positive control in the PAR binding assay. Subsequently, filters were incubated with PAR diluted in TBS-T (10 mM Tris-HCl pH 8.0, 150 mM NaCl, 0.1% Tween 20) for 1 h at room temperature. After high-stringency salt washes, protein bound PAR were detected using anti-PAR monoclonal antibody (mouse Mab ALX-804220).

### ChIP and BrdU-ChIP

ChIP was performed after double thymidine blockade and PARP1 and TRF1 RNAi. Olaparib 2 μM (AZD2281 Selleckchem) and NU1025 200 μM (Sigma Aldrich) were given to cells during release from cell cycle blockade. Cells were collected every 2 hours post-release after addition of formaldehyde (1%) directly to culture medium for 10 min at R.T. and sonicated chromatin (80 μg/sample) was immunoprecipitated (IP) overnight at 4°C with 4 μg of the anti-TRF1 antibody (goat Pab sc-1977, Santa Cruz) or the anti-WRN antibody (Rabbit Pab NB100-471, Novus Biologicals). Crosslink was then reversed with NaCl 5 M and DNA was extracted with phenol-chloroform method. Brdu-ChIP was performed after addition of 20 μM BrdU (5-bromo-2′-deoxyuridine, Sigma Aldrich) directly to HeLa culture medium for 1 h, then cells were collected and 60 μg of sonicated chromatin was incubated overnight at 4°C with 20 μl of the anti-BrdU antibody (347580 Pure BD). Then, IP was performed as described above. After precipitation with each antibody, the precipitants were blotted onto Hybond-N membrane (Amersham), and telomeric repeat sequences were detected with a Telo probe (TTAGGG). A nonspecific probe (Alu) was also used. To verify that an equivalent amount of chromatin was used in the immunoprecipitated, samples representing the 0.1%, 0.01%, and 0.001% of the total chromatin (input) were included in the dot blot. The filter was exposed to a PhosphorImager screen (Bio-Rad), and the signals were measured using ImageQuant software (Quality One; Bio-Rad).

### Electro Mobility Shift Assay (EMSA)

Telomeric duplex DNA 5′-GGGTTAGGGTTAGGGTTAGGGTTAGGGTTAGGGTTAGGGTTAGGGCCCCTC-3′ and antisense (5′-GAGGGGCCCTAACCCTAACCCTAACCCTAACCCTAACCCTAACCCTAACCC-3′ was end-labeled with [γ-^32^P]ATP (Amersham Biosciences) and T4-polynucleotide kinase (New England BioLabs) and purified from free nucleotides through G25 spin columns (GE Healthcare). Binding was carried out by incubating 0.5 ng of labelled DNA with 1 μg of unmodified or PARP1 covalently PARylated TRF1 (as above described) in 15 μl of a reaction mix of 20-mM Hepes (pH 7.9), 100-mM NaCl, 50-mM KCl, 1-mM MgCl_2_, 0.1-mM ethylenediaminetetraacetic acid (EDTA), 1 mM DTT, 5% (v/v) glycerol, 0.5 mg/ml of BSA and 0.1% (v/v) NP-40. Samples were incubated at 4°C for 90 min and then run on native 4.5% polyacrylamide gels. Gels were dried and exposed to PhosphorImager screens and acquired using ImageQuant (Bio-Rad), and the signals were measured using ImageQuant software (Quality One; Bio-Rad).

### PLA, IF-FISH and FISH in metaphase

For Proximity Ligation Assay (PLA) staining, HeLa cells, synchronized as above described, were fixed in 2% formaldehyde and permeabilized in 0.25% Triton X-100 in PBS for 5 min at room temperature at each endpoint. Then, samples were processed for immunolabeling with anti-TRF1 (rabbit Pab sc-6165, Santa Cruz) and anti-PARP1 (Mouse Mab ALX-804-211-R050, Enzo Life science) antibodies. PLA was performed by using the DUOLINK ® In situ detection reagents Red (Sigma-Aldrich) following the manufacturer’s instructions. For IF-FISH staining, cells, fixed and permeabilized as indicated above, were immunostained with mouseanti-phospho-Histone H2AX (Ser139) (clone JBW301, Merk Millipore) or anti p-S4/S8 RPA (Rabbit Pab Bethyl A300-245A) monoclonal antibodies followed by the by the anti-mouse IgG Alexa fluor 488 or anti-rabbit IgG Alexa fuor 555 secondary antibody (Cell Signaling). Then samples were re-fixed in 2% formaldehyde, dehydrated with ethanol series (70, 90, 100%), air dried, co-denaturated for 3 min at 80°C with a Cy3-labeled PNA probe, specific for telomere sequences (TelC-Cy3, Panagene, Daejon, South Korea), and incubated for 2 h in a humidified chamber at room temperature in the dark. After hybridization, slides were washed with 70% formamide, 10 mM TrisHCl pH7.2, BSA 0.1%, and then in TBS/Tween 0.08%, dehydrated with ethanol series, and finally counterstained with DAPI (0.5 μg/ml, Sigma-Aldrich) and mounting medium (Gelvatol Moviol, Sigma Aldrich). Images were captured at 63× magnification with a Leica DMIRE deconvolution microscope equipped with a Leica DFC 350FX camera and elaborated by a Leica LAS X software (Leica, Solms, Germany). This system permits to focus single planes inside the cell generating 3D high-resolution images. For telomere doublets analysis, chromosome spreads were obtained following 4 h incubation in colchicine 5 μM (Sigma-Aldrich) and prepared following standard procedure consisting of treatment with a hypotonic solution (75 mM KCl) for 20 min at 37 °C, followed by fixation in freshly prepared Carnoy solution (3:1 v/v methanol/acetic acid). Cells were then dropped onto slides, air dried, and utilized for cytogenetic analysis. Staining of centromeres and telomeres was performed as previously described [18] using the TelC-Cy3 PNA probe, and an Alexa488-labeled PNA probe specific for the human alphoid DNA sequence to mark centromeres (Cent-Alexa488) (both from Panagene, Daejon, South Korea). Metaphase images were captured using an Axio Imager M1 microscope (Zeiss, Jena, Germany) and the ISIS software (Metasystems, Milano, Italy). A total of 100 metaphases were analyzed for each sample in, at least, three independent experiments. Telomere length was calculated as the ratio between the relative fluorescence intensity of each telomere signal (T) and the relative fluorescence intensity of the centromere of chromosome 2 (C) and expressed as percentage (T/C %)[19].

## General

I would like to thank Prof. Eric Gilson and Prof. Stefano Cacchione for their generous advices and suggestions and for sharing reagents and Mr. Rocco Fraioli for technical assistance in EMSA assay.

## Funding

We gratefully acknowledge the Italian Association for Cancer Research for financial support [Grant 16910 to A.B.,17121 to E.S]. E.P. and L.P. were recipients of a fellowship from the Italian Association for Cancer Research (AIRC)

## Author contributions[Office2]

Conceptualization, S.E., S.A., A.A. G.G. and B.A.; methodology, validation, formal analysis, data curation, M.C., D.S.A., D.C., B.F., P.E., V.E., P.L., S.E.; investigation, D.S.A., M.C., D.C., P.E., V.E., P.L., B.A., and S.E.; writing original draft preparation, S.E.; writing review and editing, S.E., B.A., A.A., S.A., P.E., G.G.; visualization, S.E., B.A.; supervision, S.E., B.A.; project administration, S.E.; funding acquisition, S.E. and B.A.

## Competing interests

The authors declare no competing interests.

## Data availability statement

Raw data are available at: https://gbox.garr.it/garrbox/index.php/s/LJjfkvEA0tvYsf8

---

**Fig. S1.**
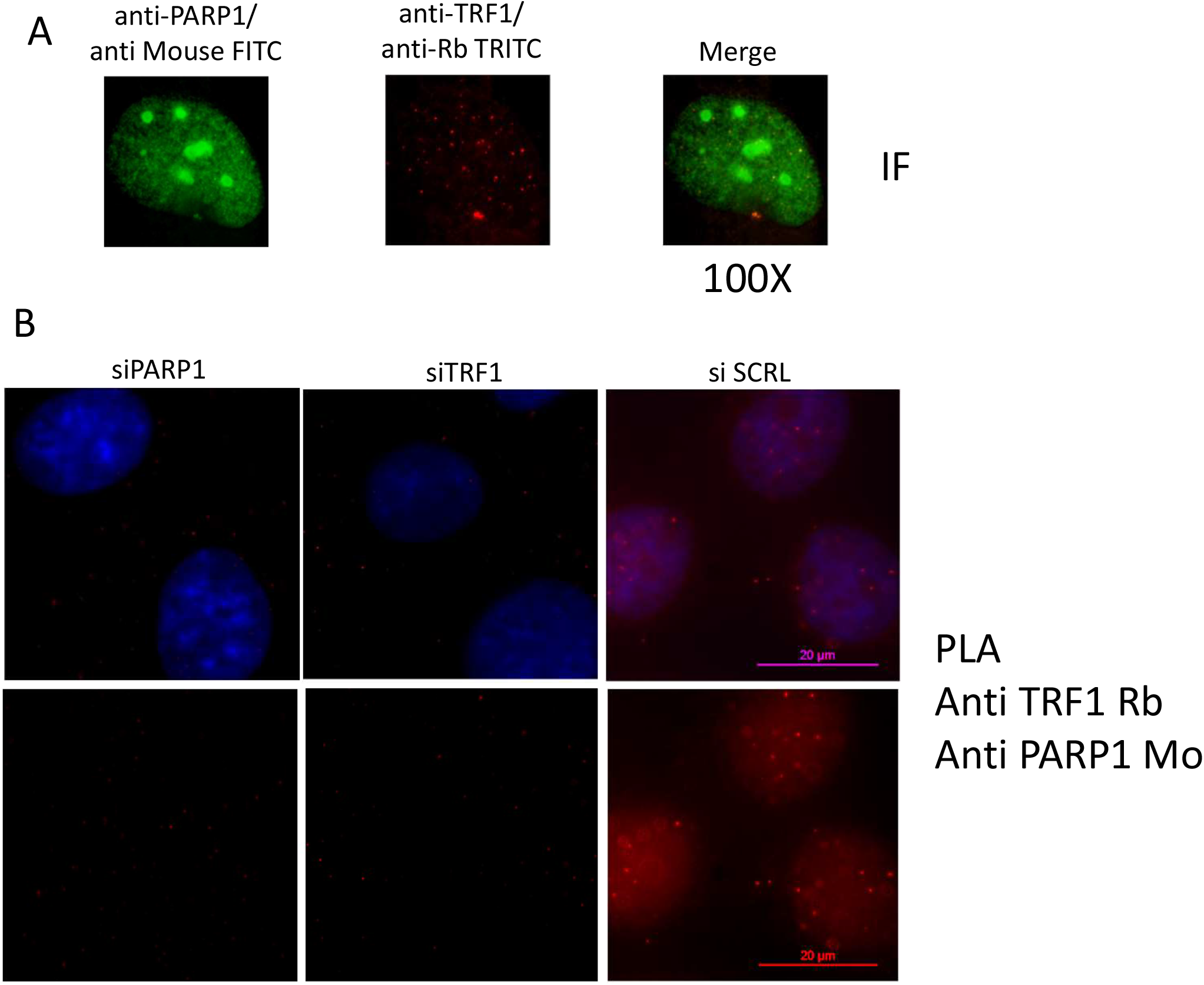
A: Hela cells were seeded, fixed and processed for co-immunofluorescence with anti-mouse PARP1 and anti-rabbitTRF1 specific antibodies followed by the indicated secondary antibody. Fluorescent signals corresponding to both protein staining are shown in representative images at 100X magnification. B: cells transfected with the indicated siRNAs were processed for co-IF with the above primary antibodies and processed for Proximity Ligation Assay with the DUOLink Red kit mouse/rabbit (Sigma). Representative images at 63X magnification are shown.

**Fig. S2.**
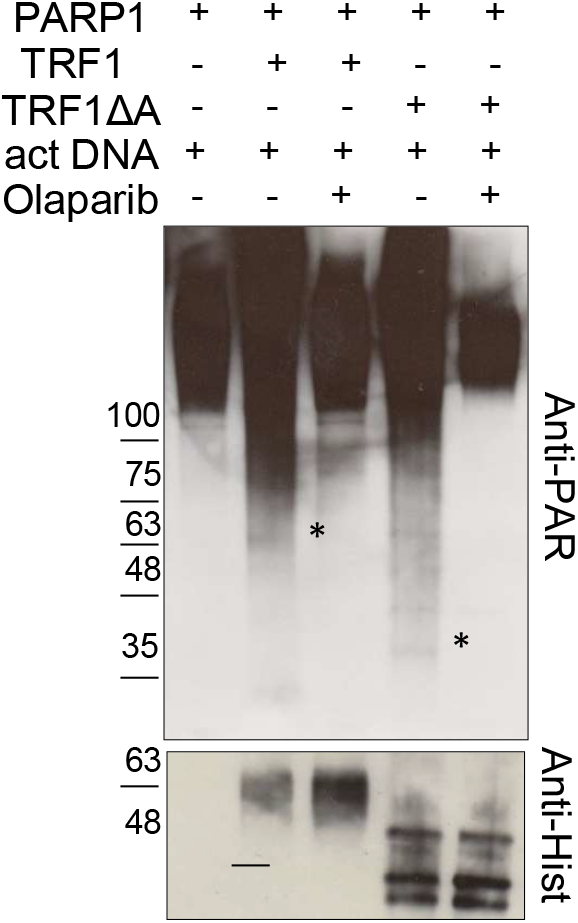
high activity purified PARP1 enzyme was incubated with NAD+ in the PARylation reaction buffer in absence or presence of recombinant His-tag full length TRF1 or delta acidic TRF1 (ΔA), with or without activating DNA or the PARP1 inhibitor Olaparib, 5 μM (**B**). Protein mixtures were resolved on PAGE and decorated with an anti-PAR specific antibody or anti-His antibody to detect TRF1 isoforms where present..

**Fig. S3.**
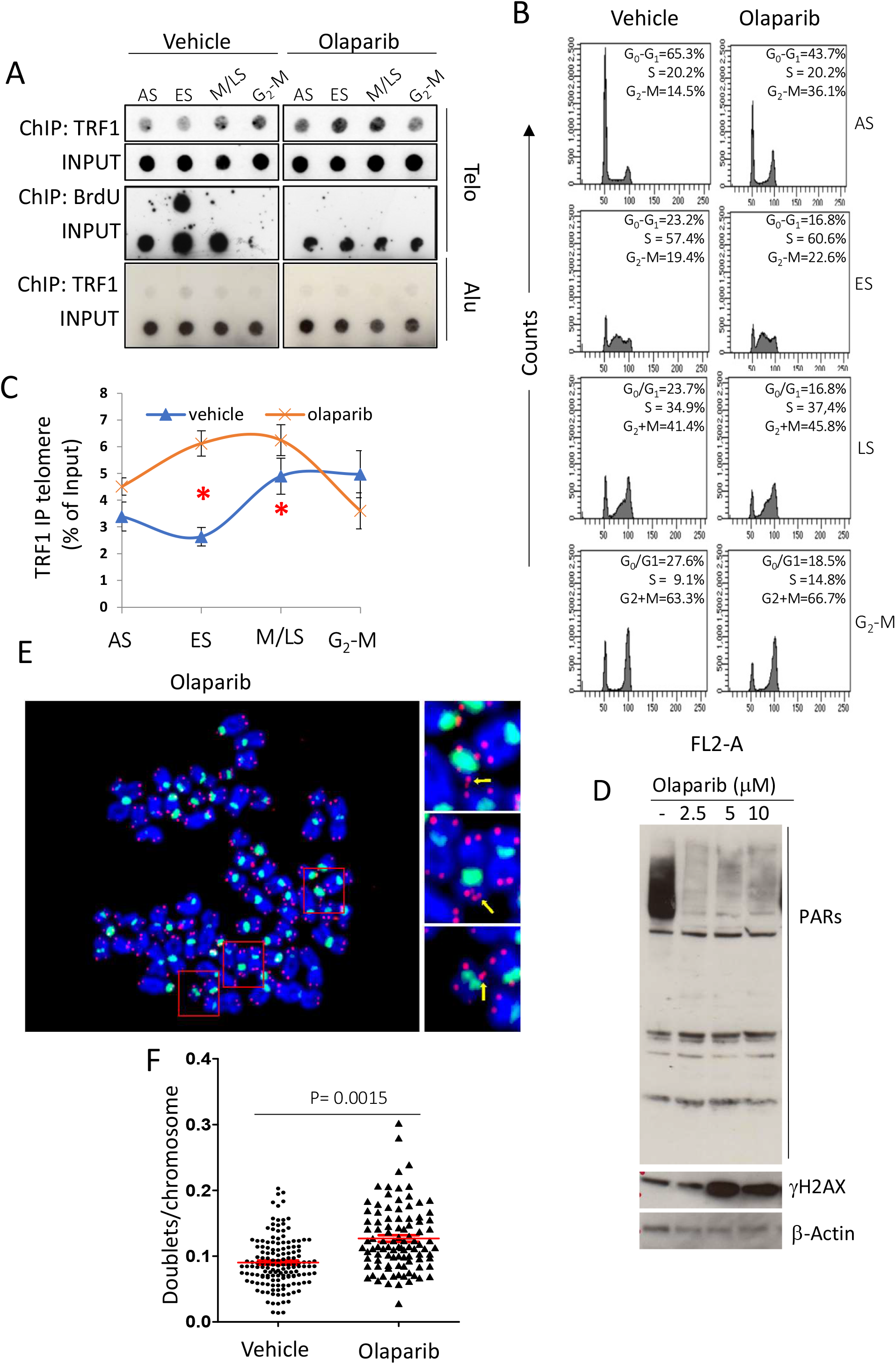
Olaparib perturbs telomere replication and induce telomere fragility. Hela cells were synchronized in the early S phase by double thymidine block and then released. 2 μM Olaparib was administered where indicated, from the second thymidine block. 1 hr pulse of BrdU incorporation was performed before sample collection at the indicated time points. Samples collected underwent ChIP with an anti-TRF1 specific antibody (**A**) or flow cytometry to control cell cycle distribution (**B**). Immunoprecipitated chromatin samples were dot blotted and processed first by western blot against BrdU, and then hybridized with ^32^P labelled telo or alu probes. Signals were quantified by densitometry and reported in graphs as the percentage of each relative input after normalization on the Alu signals. Stars indicate BrdU incorporation. (**C). D**: Hela cells were treated with the indicated doses of Olaparib. Then cells were collected and processed for western blot analysis against anti-PAR antibody and γ-H2AX. β-actin was used as a loading control. **E**: representative image at 100X magnification of pantelomeric/pancentromeric FISH analysis of Hela metaphase spread after exposure to 2 μM Olaparib for 72 hours showing doublets formation. The number of doublets for chromosome was scored and reported in graphs in **F**.

**Figure S4.**
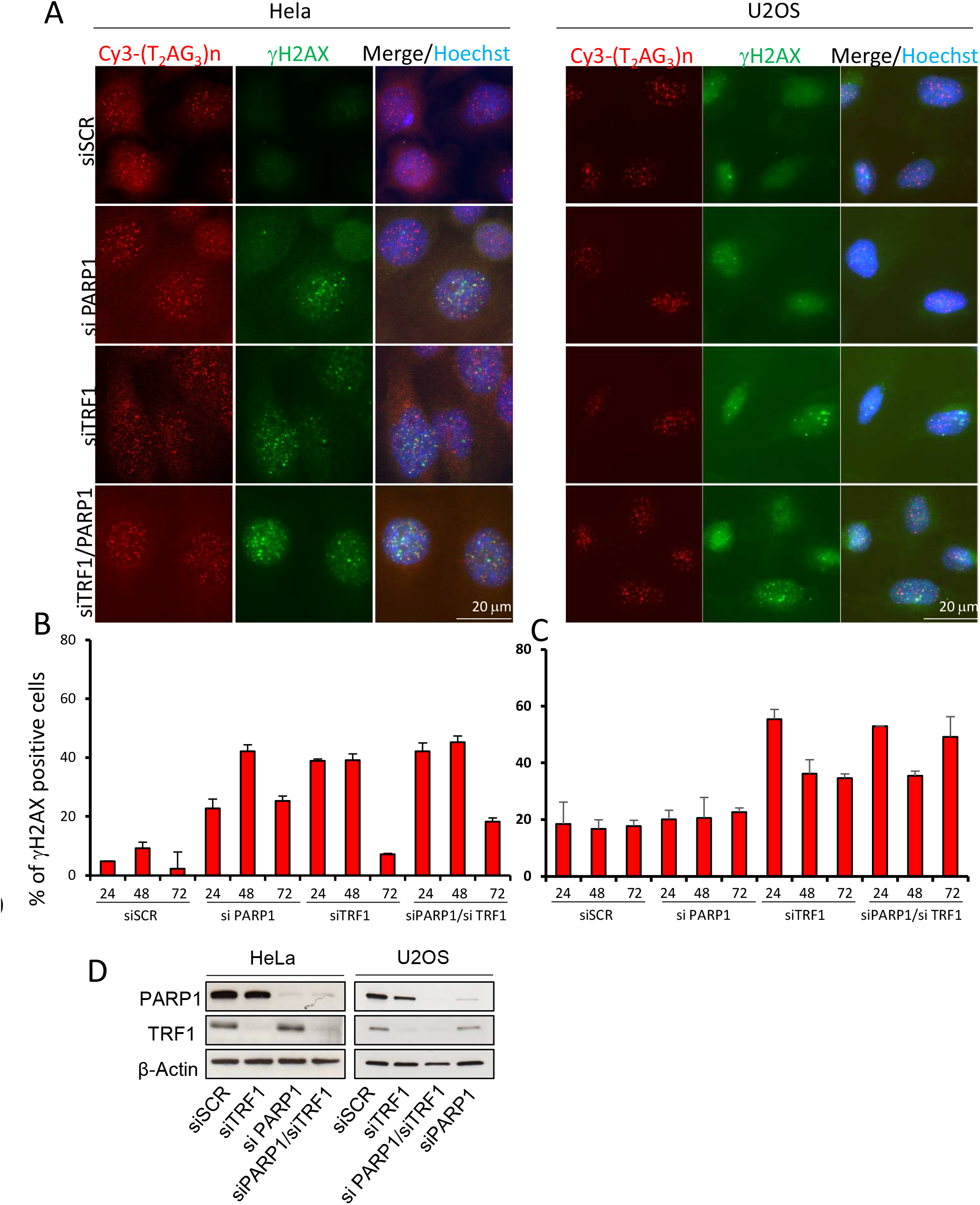
siPARP1 induces DDR in Hela but not in U2OS and its effect is epistatic to TRF1. Hela and U2OS cells were transfected with siRNAs against siPARP1 and siTRF1 alone and in combination and against a scrambled sequence. Then samples were fixed at the indicated endpoints after transfection and processed for IF-FISH against γH2AX and telomere repeats with a Cy3-Telo PNA probe and counterstained with DAPI. Signals were acquired by Leica Deconvolution fluorescence microscope at 63X magnification (representative images are shown in panel **A**). The percentage of γH2AX positive cells in Hela and U2OS was scored and reported in histograms in **B** and **C** respectively. **D** western blot of interfered cells for the control of protein depletion. The mean of three independent experiments is shown for each sample. Bars are SD.

**Fig S5.**
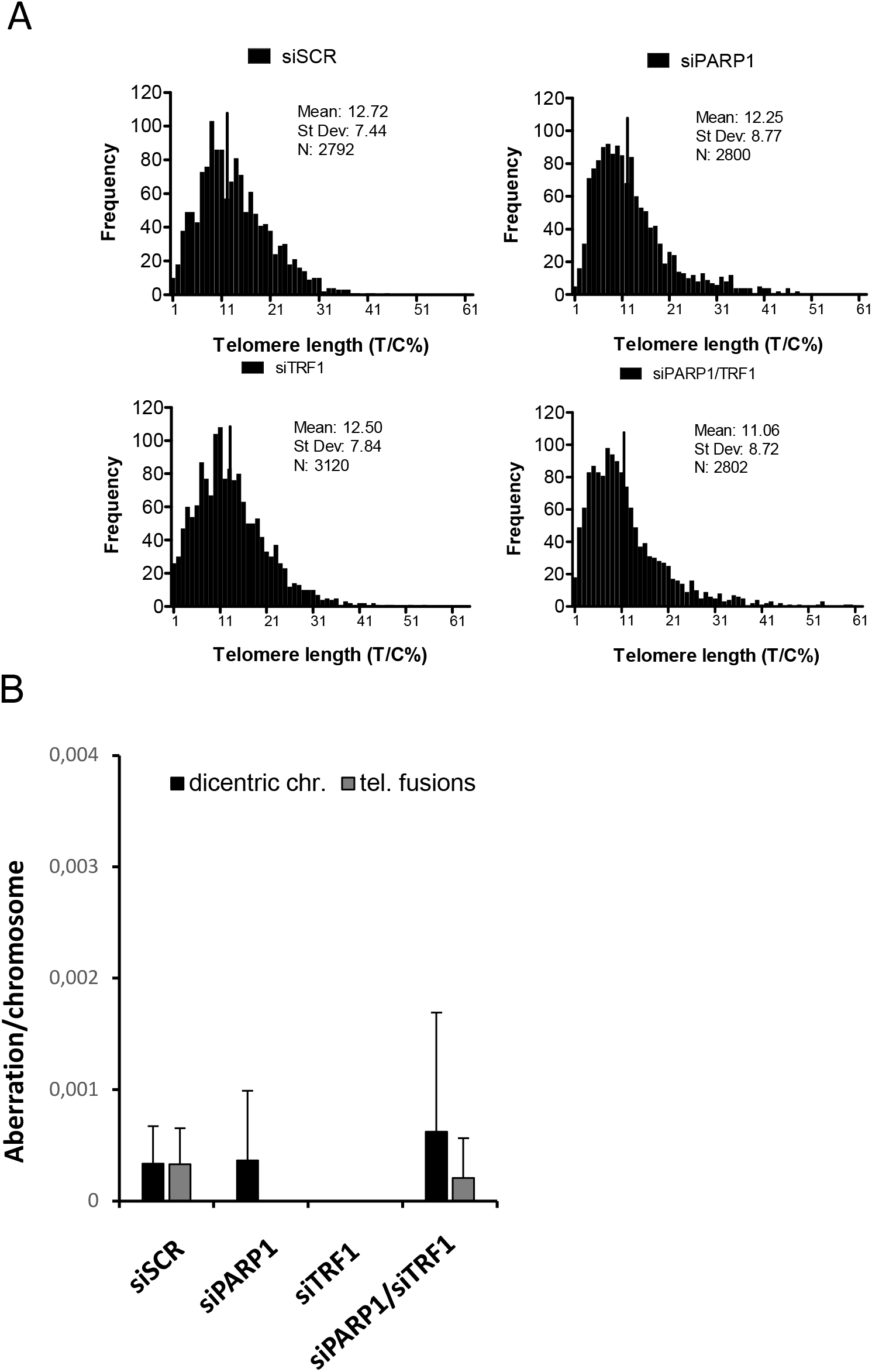
Effect of TRF1/PARP1 interference on telomere length and other telomere aberrations. Hela cells interfered for 72 hours with the indicated siRNAs were synchronized in metaphase and processed for FISH analysis with pan-centromeric and pan-telomeric probes. Images of metaphases were acquired and analyzed for telomere length (**A**) and scored for the presence of dicentric chormosomes and telomeric fusions (**B**). As shown by the histograms, both telomere length and the number of aberrations/chromosome were not affected by transfection. (test t student p> 0,1)

**Fig S6.**
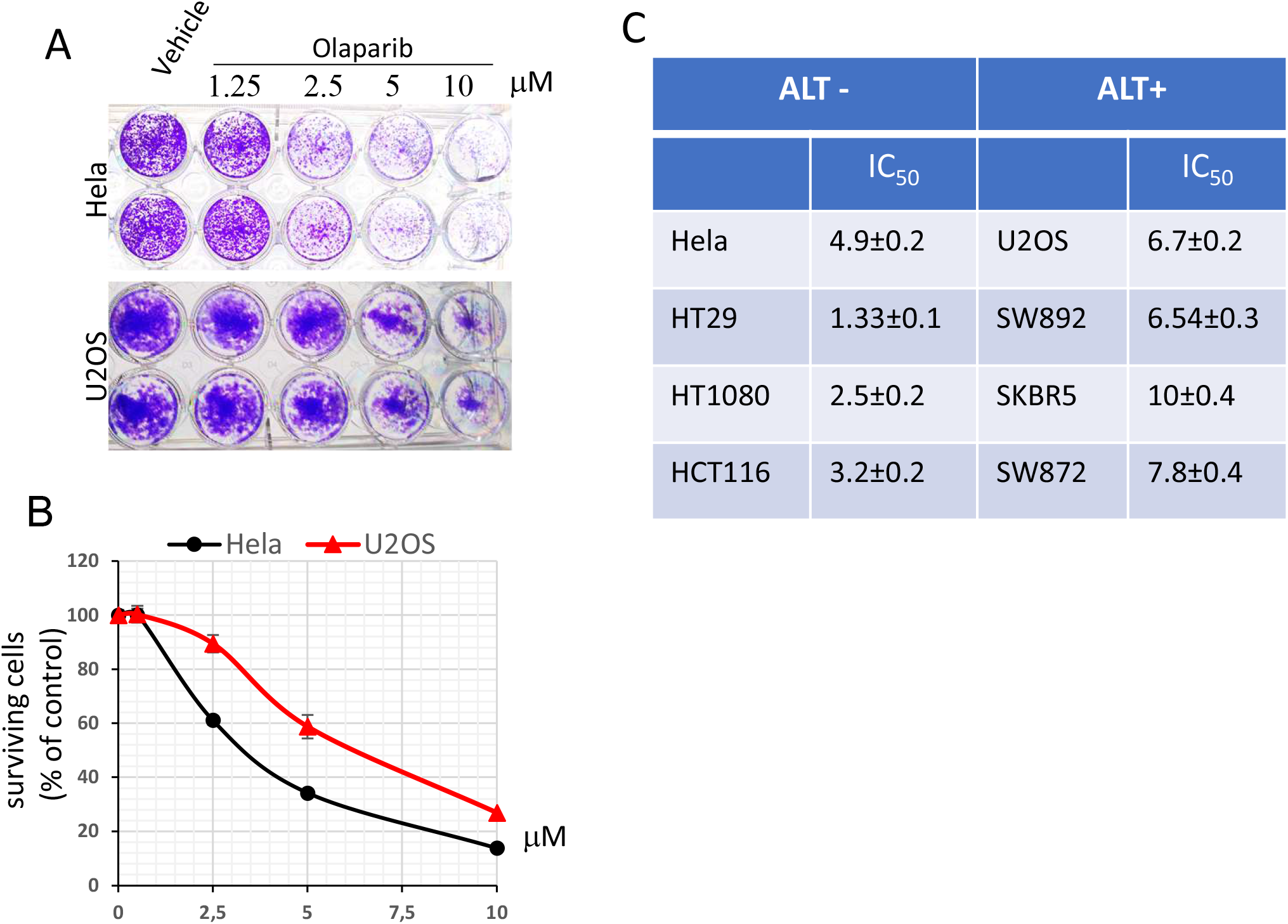
Differential effect of Olaparib in ALT and non-ALT cells. **A:** Hela and U2OS cells were seeded and exposed to the indicated doses of Olaparib for 7 days. Then cells were fixed and stained with crystal violet. Representative images are shown. **B:** Samples in **A** were quantified by spectrophotometric analysis to determine cell survival. Graphs represent the percentage of surviving cells with respect to DMSO treated samples. **C**. The indicated cell lines underwent Olaparib treatment and crystal violet as above described. IC_50_ values were calculated and reported ± standard deviation.

## Notes

### Competing Interest Statement

The authors have declared no competing interest.

